# The GMD-biplot and its application to microbiome data

**DOI:** 10.1101/814269

**Authors:** Yue Wang, Timothy W Randolph, Ali Shojaie, Jing Ma

## Abstract

Exploratory analysis of human microbiome data is often based on dimension-reduced graphical displays derived from similarities based on non-Euclidean distances, such as UniFrac or Bray-Curtis. However, a display of this type, often referred to as the principal coordinate analysis (PCoA) plot, does not reveal which taxa are related to the observed clustering because the configuration of samples is not based on a coordinate system in which both the samples and variables can be represented. The reason is that the PCoA plot is based on the eigen-decomposition of a similarity matrix and not the singular value decomposition (SVD) of the sample-by-abundance matrix. We propose a novel biplot that is based on an extension of the SVD, called the generalized matrix decomposition (GMD), which involves an arbitrary matrix of similarities and the original matrix of variable measures, such as taxon abundances. As in a traditional biplot, points represent the samples and arrows represent the variables. The proposed GMD-biplot is illustrated by analyzing multiple real and simulated data sets which demonstrate that the GMD-biplot provides improved clustering capability and a more meaningful relationship between the arrows and the points.

## Introduction

A biplot simultaneously displays, in two dimensions, rows (samples) of a data matrix as points, and columns (variables) as arrows. Based on a matrix decomposition of the data matrix, the biplot is a useful graphical tool for visualizing the structure of large data matrices. It displays a dimension-reduced configuration of samples, as in a PCoA plot, and the variables with respect to the same set of coordinates. If meaningful sample groupings are observed, this allows for visualizing which variables contribute most to the separation. The traditional biplot, as first introduced in [1], displays the first two left and right singular vectors of the singular value decomposition (SVD) of the data matrix as points and arrows, respectively. This biplot, which we hereafter refer to as the SVD-biplot, uses the SVD to find the optimal least-square representation of the data matrix in a low-dimensional space. The SVD-biplot can show Euclidean distances between samples and also display approximate variances and correlations of the variables. It also has the appealing property that the singular values obtained from the SVD are non-increasing, indicating that the decomposition of the total variance of the data matrix into each dimension is non-increasing.

In many scenarios, the Euclidean distance may not be optimal for characterizing dissimilarities between samples. An important example arises in the analysis of microbiome data, in which marker gene sequences (e.g., 16s rRNA) are often grouped into taxonomic categories using bioinformatics pipelines such as QIIME [2] or Mothur [3]. These taxon counts can be summarized into a data matrix with rows and columns representing samples and taxon abundances, respectively. A variety of non-Euclidean distance measures, including nonlinear measures, are then used to quantify the similarity between samples. One common measure of dissimilarity is the UniFrac distance (weighted or unweighted), which is a function of the phylogenetic dissimilarity of a pair of samples [4; 5]. Other non-phylogenetic, non-Euclidean dissimilarities include Jaccard or Bray-Curtis distances (see, e.g., [6] and the references therein). Plotting the samples in the space of the first few principal components (PCs) of the similarity matrix obtained from such non-Euclidean distance matrices— often referred to as principal co-ordinates analysis (PCoA)—may reveal an informative separation between samples. However, the configuration of samples yielded by PCoA only keeps pairwise distances between samples and lacks a coordinate system that relates to the taxa which constitute each sample. Hence, it does not shed any light on which taxa may play a role in this separation. One approach for addressing this problem is to simply plot an arrow for each taxon based on its correlation with the first two PCs of the non-Euclidean similarity matrix [7]. However, in such a “joint plot” [8], the direction and length of an arrow does not represent the taxon’s true contribution to the dissimilarity between samples. In addition, due to the lack of a coordinate system, one cannot add sample points for future observations into this “joint plot”.

Three main approaches have been recently proposed to extend the SVD-biplot to more general distances defined on the samples. The R package “ade4” [9] provides a biplot that can handle weighted Euclidean distances but cannot apply to non-Euclidean distances. The second approach, proposed by [10], aims to approximate the non-Euclidean distance by a weighted Euclidean distance. Weights are estimated for variables and the biplot can subsequently be constructed using weighted least-square approximation of the matrix. This approach has a straightforward interpretation. However, the estimated weighted Euclidean distance may not capture all the information from the original non-Euclidean distance. A recent proposal in [11] appears to be the first to address the lack of mathematical duality between the samples’ locations (points) and the variables’ contribution (arrows) to those locations. This approach seeks an approximate SVD-like decomposition of the data matrix, which directly takes the non-Euclidean distance into consideration. This SVD-like decomposition has the following two advantages. First, the left singular vectors are the eigenvectors of the similarity measure derived from the non-Euclidean distance, which preserve the role of the non-Euclidean distance in classifying the samples. Second, an approximate matrix duality (AMD) between the left and right singular vectors is restored, which simply means that each set of vectors can be immediately obtained from the other. To emphasize this connection, we hereafter refer to this decomposition as the AMD. Unfortunately, the AMD also suffers from two drawbacks. First, the AMD is only an approximate decomposition of the data matrix, and hence may not capture all the variation of the original data. In particular, the configuration of samples displayed in an AMD-biplot is independent of the data matrix, since the left singular vectors of the AMD only depend on the non-Euclidean distances. Ignoring the data matrix for classifying samples seems non-intuitive since the data matrix is typically assumed to contain some information on the sample similarities. Second, the AMD may result in non-decreasing “singular values”. While these seem like minor technical issues, the second drawback can have important practical implications: which of the left and right singular vectors should be displayed in the resulting biplot? The authors of [11] suggest constructing the AMD-biplot based on the two left and right singular vectors that correspond to the two largest singular values. This AMD-biplot assures that the arrows for variables are as meaningful as possible, but may fail to reveal meaningful sample clusters if the information of sample clusters is only associated with the first several left singular vectors. An alternative approach may be to simply display the first and second left and right singular vectors of the AMD (as done for the SVD). Unfortunately, this strategy does not solve the problem either: although we may observe meaningful sample clusters, the arrows may not be meaningful due to the small singular values. There is thus a lack of clarity regarding which singular vectors should be used to construct the AMD-biplot.

The drawbacks of the AMD motivate our proposal which is based on the generalized matrix decomposition (GMD) [12]. The GMD is a direct generalization of the SVD that accounts for structural dependencies among the samples and/or variables. This approach has several advantages. First, as with the AMD, it directly handles any non-Euclidean distance matrix. Specifically, the full information from that distance matrix is used. Second, unlike the AMD, which provides an approximate decomposition of the data matrix, the GMD provides an exact decomposition of the original data matrix without losing any information. Third, the GMD restores the matrix duality in a mathematically rigorous manner, unlike the approximate matrix duality obtained with the AMD; it naturally extends the duality inherent in the SVD and allows one to plot both the configuration of samples and the contribution of individual variables with respect to a new coordinate system. Fourth, the GMD gives non-increasing GMD values and so the resulting GMD-biplot can be directly constructed based on the first two left and right GMD vectors. Lastly, unlike the AMD-biplot whose sample clusters only depend on the distance, the GMD-biplot uses both the non-Euclidean distance and the data matrix for classifying samples, which more directly connects the contribution of the individual variables to the configuration of samples. Additionally, besides accounting for the non-Euclidean distances between samples, the GMD can also account for auxiliary information on (dis)similarities between the variables.

In the following, we first summarize the GMD-biplot framework and then compare the GMD, AMD and SVD biplots in three numerical studies. We then discuss advantages of the proposed GMD-biplot and further extensions.

## Materials and Methods

We denote the data matrix by **X** ∈ ℝ^*n*×*p*^, where *n* is the number of samples and *p* is the number of variables (taxa). We assume that the columns of **X** are centered to have mean 0 and rank(**X**) = *K* ≤ min(*n, p*). For any matrix **M**, we denote its *i*^*th*^ row (sample) and its (*i, j*) entry by **m**_*i*_ and *m*_*ij*_, respectively. We denote the transpose of **M** by **M**^*T*^.

### Biplot, distance measure and the AMD

A biplot is a graphical method to simultaneously represent, in two dimensions, both the rows (as points) and columns (as arrows) of the matrix **X** on the same coordinate axes. Given a decomposition of **X** as **X** = **AB**^*T*^, a biplot displays two selected columns of **A** and **B**. The SVD-biplot is based on the SVD of **X**, i.e. **X** = **USV**^*T*^, where **U**^*T*^ **U** = **I**_*K*_, **V**^*T*^ **V** = **I**_*K*_ and **S** = diag(*σ*_1_, …, *σ*_*K*_) with *σ*_1_, …, *σ*_*K*_ being a sequence of non-increasing and positive singular values. Here **I**_*K*_ is a rank *K* identity matrix. Based on the SVD, **A** and **B** can take various forms; examples include form and covariance biplots [7]. Since our primary interest is to visualize the clustering of samples, we focus on the form biplot in this paper and comment on the covariance biplot in the Discussion.

The SVD-biplot displays the first two columns of **US** and **V**, which can explain 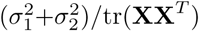 of the total variance of **X**. The SVD of **X** is closely related to the eigen-decomposition of the similarity kernel **XX**^*T*^, as we can write **XX**^*T*^ = **US**^2^**U**^*T*^. Thus, the eigen-decomposition of **XX**^*T*^ provides a way to calculate **U** and **S**. Once **U** and **S** are calculated, one can calculate **V** from the duality between **U** and **V**; that is, **VS** = **X**^*T*^ **U**. The similarity kernel **XX**^*T*^ characterizes the Euclidean distance between samples. To see this, we define the Euclidean squared distance between the *i*^*th*^ and *j*^*th*^ sample as 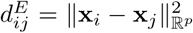. Let 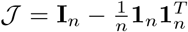 be the centering matrix where **1**_*n*_ is an *n* × 1 vector of ones. It can then be seen that 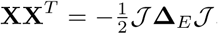, where the (*i, j*) entry of **Δ**_*E*_ is 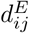.

Now, if **Δ**_*E*_ is replaced by a matrix **D** of non-Euclidean squared dissimilarities, one can still define a similarity kernel by 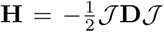. One such example is when **D** arises from distances between sample vectors of microbial abundances (or presence/absence) which account for a phylogenetic tree, as in a weighted (respectively, unweighted) UniFrac distance matrix. In this case, one can construct a principal coordinate analysis (PCoA) plot of the samples using 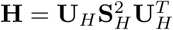. However, an SVD-biplot cannot be constructed, since there is no **V** that corresponds to the variables. The AMD addresses this problem by fixing **U**_*H*_ and then seeking a matrix **V**_*H*_ with orthonormal columns and a diagonal matrix **D**_*H*_ with non-negative elements that minimize the objective function

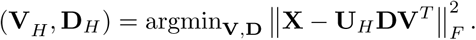

Here, 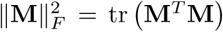 is the Frobenius norm of **M**, and for any square matrix **M** ∈ ℝ^*d*×*d*^, 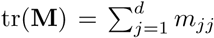. The resulting AMD-biplot can be constructed by plotting the two columns of **U**_*H*_ **D**_*H*_, as sample points, and **V**_*H*_, as arrows for variables; the selected two columns/rows correspond to the two largest elements of **D**_*H*_.

### GMD and the GMD-biplot

The concept of the generalized matrix decomposition (GMD) was introduced by Escoufier [13] and further developed in [12]. It is a generalization of the SVD with additional structural dependencies taken into consideration. We briefly review the key ideas behind the GMD. Let **H** ∈ ℝ^*n*×*n*^ and **R** ∈ ℝ^*p*×*p*^ be two positive semi-definite matrices, which, respectively, characterize the similarities between samples and between variables. The **H, R**-norm of **X** is defined as 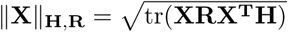. For any *q* ≤ *K*, the GMD solution 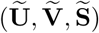 finds the best rank-*q* approximation to **X** with respect to the **H, R**-norm, that is,

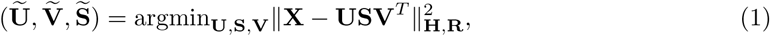

subject to **U**^*T*^ **HU** = **I**_*q*_, **V**^*T*^ **RV** = **I**_*q*_ and diag(**S**) *≥* 0. Here, **Ũ** and 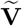 are the left and right GMD vectors, respectively, and 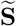 is a diagonal matrix containing the GMD values. Note that **Ũ** and 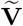 are orthogonal with respect to **H** and **R** respectively, but they may not be orthogonal with respect to the Euclidean norm unless **H** = **I**_*n*_ and **R** = **I**_*p*_. The following property of the GMD provides a way to calculate the GMD components; the proof can be found in [13].

#### Proposition 1

The GMD solutions 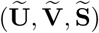 satisfy:

a. 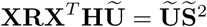
b. 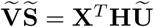

Proposition 1(a) suggests that the diagonal elements of 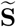 and corresponding columns of **Ũ** are eigenvalues and corresponding eigenvectors of **XRX**^*T*^ **H** respectively. Proposition 1(b) establishes the duality between **Ũ** and 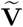, meaning that 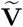 can be immediately obtained given **Ũ** and 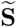. Alternatively, an efficient algorithm for finding the solution to Eq. (1) was proposed in [12], which is less computationally intensive compared to finding the eigenvalues and eigenvectors of **XRX**^*T*^ **H**. The algorithm also ensures that the diagonal elements of 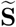 are non-increasing.

Note that the GMD can handle the non-Euclidean similarity kernel **H** just by taking **R** = **I**_*p*_. Based on the GMD of **X** with respect to **H**, the GMD-biplot can be constructed with respect to the coordinate system provided by the first two columns of **V**. More specifically, letting **v**_*j*_ be the *j*-th column of **V**, the *i*-th sample point can be configured by the coordinates of **x**_*i*_, given by 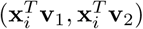. To plot the arrow for the *j*-th variable, we consider the vector **e**_*j*_ ∈ ℝ^*p*^, which has a 1 in the *j*-th element and 0’s elsewhere. Then, the arrow for the *j*-th variable can be configured by the coordinates of **e**_*j*_, given by 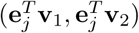. This coordinate system also allows the configuration of future samples. Letting **x**_*_ ∈ ℝ^*p*^ be a future sample, we can add **x**_*_ into the GMD-biplot as a point located at 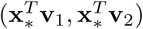. Similar to the SVD-biplot, the GMD-biplot can explain 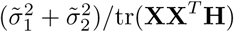 of the total variance of **X** with respect to the **H, I**_*p*_ norm, where 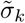 is the *k*-th diagonal element of 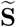 for *k* = 1, 2.

Since the GMD values are non-increasing, for the purpose of constructing the GMD-biplot, we can choose *q* = 2 in the GMD problem (Eq. (1)), which may save considerable computational time. In contrast, since the AMD may produce non-decreasing “singular values”, we have to find the full decomposition of **X** by the AMD before deciding which singular vectors to plot in the AMD-biplot; this may become computationally intensive for large *n* and *p*.

## Results

In the results below, we compare the GMD, AMD and SVD biplots on three data sets in the manner that each has been proposed recently for microbiome data. In particular, in [11], the AMD-biplot is advocated specifically for relative abundance data, while in [14] the SVD-biplot is advocated for data that have been scaled by the centered log-ratio (CLR) transformation. The GMD-biplot is constructed using the CLR-transformed data. We first examine the performance of all biplots using the smokeless tobacco data set explored in [11]. In the second study, we compare their performances using the human gut microbiome data from [15]. In the third analysis, we simulate a data set based on the smokeless tobacco data to illustrate a dilemma that the AMD-biplot may face.

### Analysis of the smokeless tobacco data

This data set includes 15 smokeless tobacco products: 6 dry snuffs, 7 moist snuffs, and 2 toombak samples from Sudan. Three separate (replicate) observations (starting with sample preparation) were made of each product, so that in total 45 observations are available. Each observation has a 271 × 1 vector of taxon counts, and thus the data set can be formed into a 45 × 271 matrix, denoted by **X**. The squared weighted UniFrac distance, denoted **Δ** ∈ ℝ^45×45^, was used to measure the distance between samples. The corresponding similarity kernel **H** was calculated as 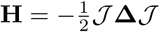. Since **H** is not positive semi-definite, we enforced it to be positive semi-definite by removing its negative eigenvalues and corresponding eigenvectors. The resulting similarity kernel, denoted **H**^*^, has rank 27.

For the GMD-biplot, we consider the CLR transformation of **X**. Specifically, denoting the geometric mean of a vector **z** by 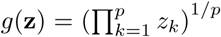, the CLR transformation of **x**; *i* = 1, …, 45 is given by

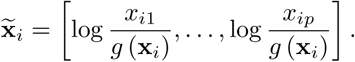

We denote the resulting data matrix by 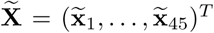. For the AMD-biplot, we converted each row of **X** into the empirical frequencies, and further centered the rows and columns to have mean 0, as done in [11]. We denote the resulting data matrix by 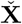.

We constructed the GMD-biplot and the AMD-biplot based on **H**^*^ using 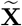 and 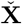, respectively. Fig. 1(d) displays the proportion of variance captured by each GMD component. It can be seen that the first two GMD components capture more than 80% of the total variance of **X**, which assures that the resulting GMD-biplot (Fig. 1(a)) visualizes the data well. As shown in Fig. 1(a), the GMD-biplot is perfectly successful at separating the different tobacco products (dry, moist and toombak). Furthermore, the replicates corresponding to the same product are tightly clustered. By examining the arrows for taxa in Fig. 1(a), we see that moist samples may be characterized by elevated levels of *alloiococcus* and *halophilus*, while *aerococcaceae* appears elevated in toomback samples.

**Figure 1:**
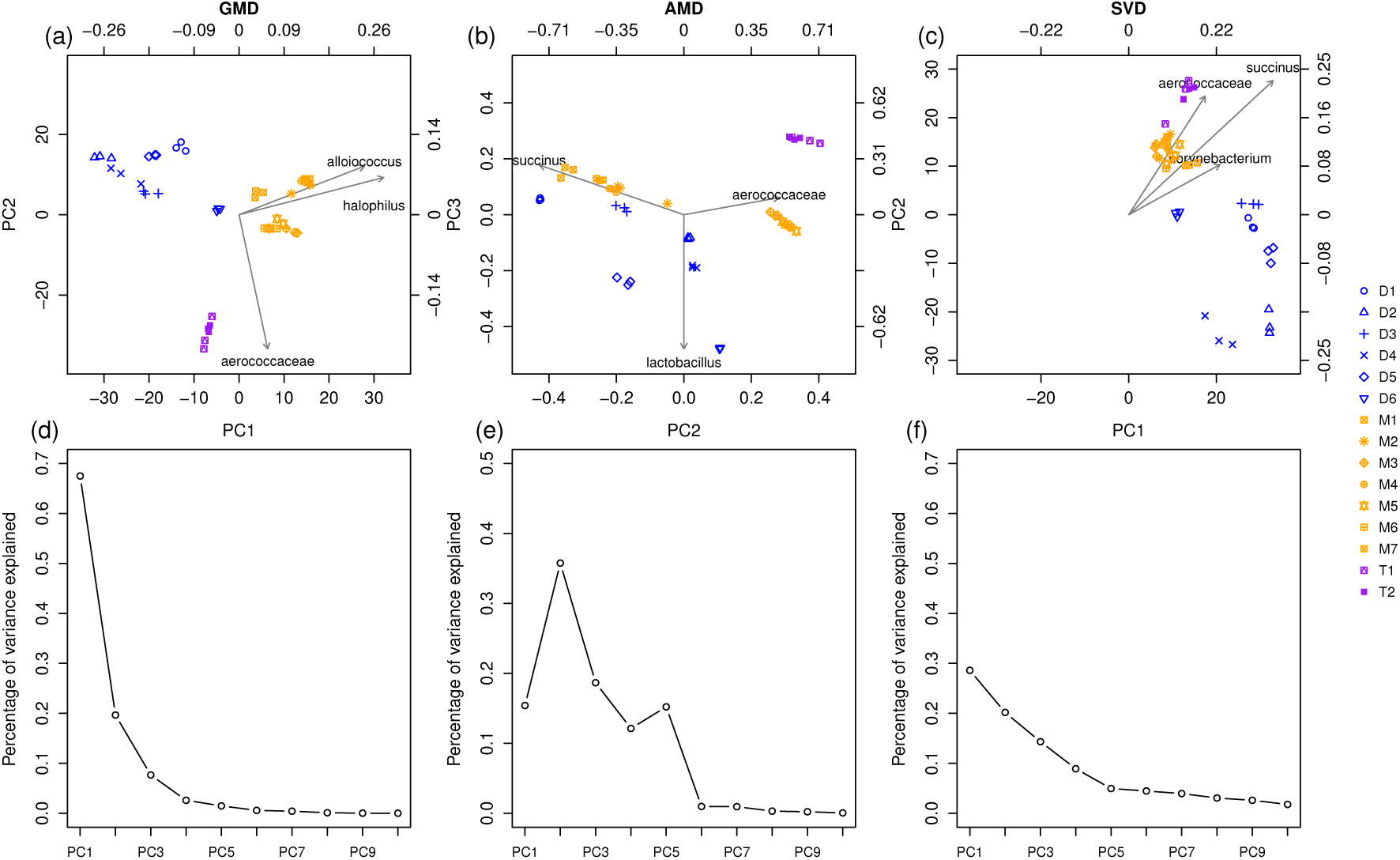
Biplots and scree plots for the analysis of smokeless tobacco data. (a): The GMD-biplot based on the first and second component; (b): The AMD-biplot based on the second and the third components; (c): The SVD-biplot based on the first and second component; (d): The GMD scree plot; (e): The AMD scree plot; (f) The SVD scree plot. All biplots (a), (b) and (c) display the top 3 taxa with the longest arrows. The samples points are colored by type (“blue” = “dry”, “orange” = “moist”, “purple” = “toombak”) and samples corresponding to replicates of the same product are plotted with the same symbol. All scree plots (d), (e) and (f) display the contribution of the top 10 components.

Fig. 1(e), which is the same as the right bottom panel of Fig. 1 in [11], shows that the AMD singular values are not necessarily decreasing. It should be noted that Fig. 1(b) is slightly different from Fig. 3 in [11]; this difference may be due to the use of **H**^*^ here as opposed to **H** in [11]. This is because we wanted the the AMD-biplot to be directly comparable to the GMD-biplot since the GMD requires both **H** and **R** to be positive semi-definite. From Fig. 1(b), it can be seen that the AMD successfully separates toombak samples (purple points) from dry (blue) and moist (orange) snuffs, although the separation between dry and moist snuffs in the AMD-biplot is not as definitive as that in the GMD-biplot (Fig. 1(a)).

**Figure 2:**
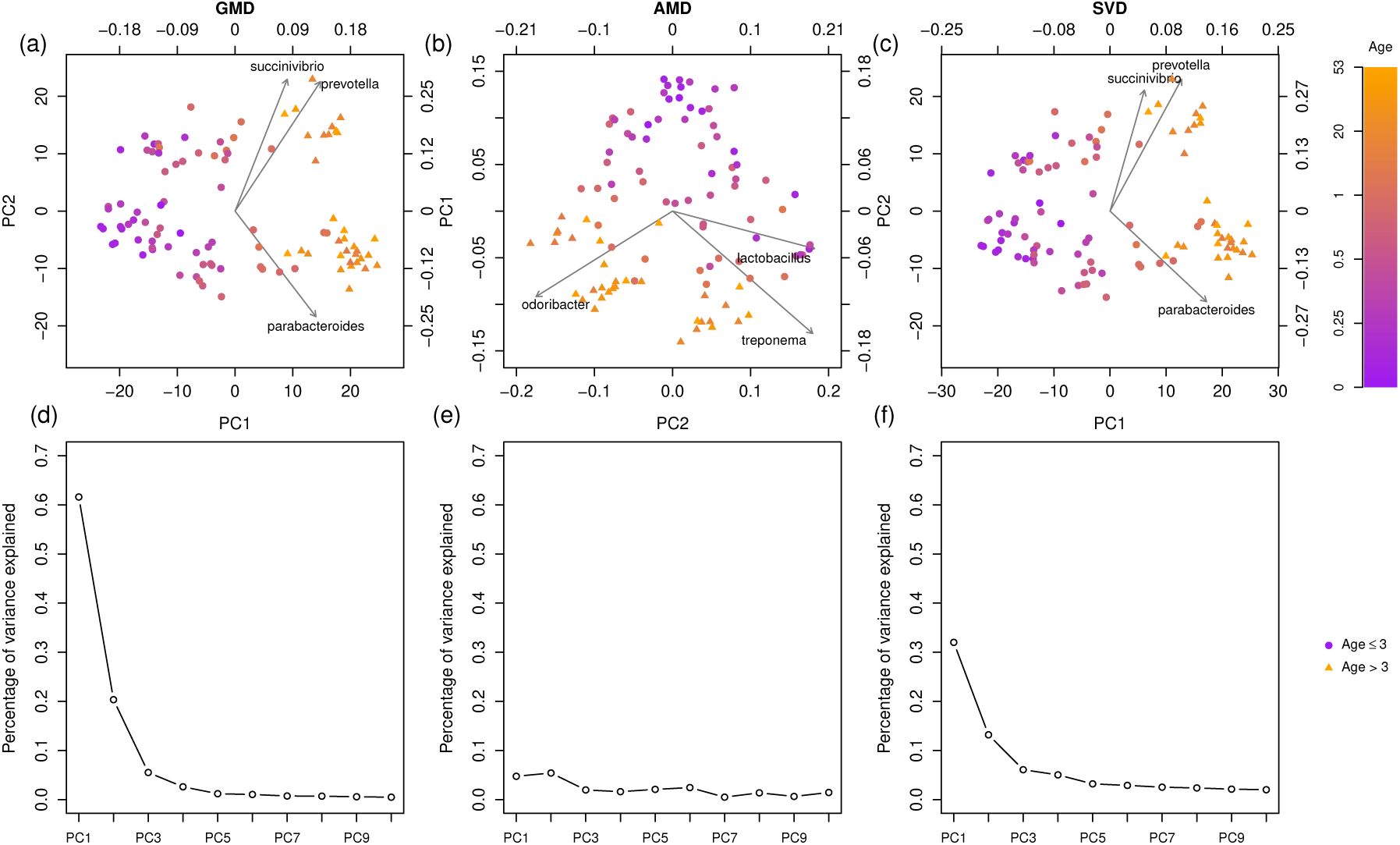
Biplots and scree plots for the analysis of the human gut microbiome data. (a): The GMD-biplot based on the first and second GMD components; (b): The AMD-biplot based on the first the second components; (c): The SVD-biplot based on the first and second SVD components; (d): The GMD scree plot; (e): The AMD scree plot; (f): The SVD scree plot. Biplots (a), (b) and (c) display the top 3 taxa with the longest arrows. Symbols of samples points are based on the ages of samples (“age ≤ 3” = “circle”; “age *>* 3” = “triangle”). Scree plots (d), (e) and (f) display the contribution of the top 10 components.

**Figure 3:**
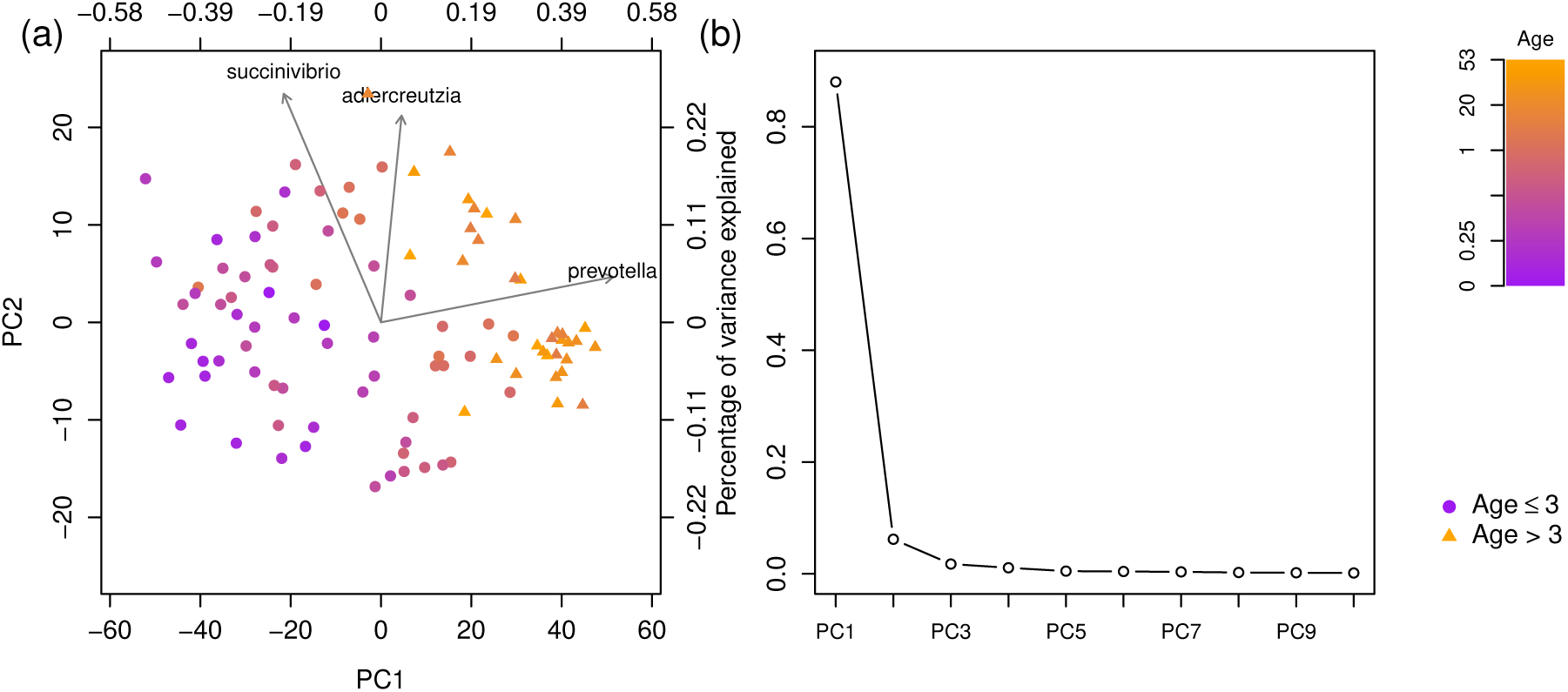
The biplot and scree plot for the analysis of the human gut microbiome data using both H and R. (a): The GMD-biplot using both **H** and **R** based on the first and second GMD components. Top 3 taxa with the longest arrows are displayed. Symbols of samples points are based on ages of samples (“age ≤ 3” = “circle”; “age *>* 3” = “triangle”) (b): The GMD scree plot using both **H** and **R**: the contributions of top 10 components are displayed.

Additionally, we included the SVD-biplot and its corresponding scree plot in Fig.1 (c) and (f) respectively. As the SVD-biplot assumes the Euclidean distance between samples, it is more appropriate to construct the SVD-biplot using the CLR transformed data 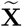 than the relative abundance data 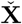 [14]. It can be seen from Fig. 1(c) that although the SVD successfully separates dry snuffs from moist and toombak samples, it does not give a clear separation between moist snuffs and toombak samples.

It is worth noting that the three biplots idenfity different top taxa, i.e, the taxa with the longest arrows. Although a biplot is not a rigorous statistical method to detect important taxa, it may shed light on which taxa are important to the observed sample clustering. To see this, we performed a univariate linear regression of each taxon (each column of 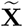) on the tobacco groups (dry, moist and toombak), and obtained *p*−values representing the strength of association between each taxon and the tobacco groups. We then sorted these *p*−values in a non-decreasing order, and obtained the rank of each taxon based on the sorted *p*−values. Hence, it is desirable that the taxa with the lowest ranks can be identified by the biplots. Table S1 summarizes the ranks of the top 10 taxa identified by each biplot. It can be seen that the top 10 taxa identified by the GMD-biplot have lower ranks on average than those identified by the AMD and SVD biplots, indicating that the GMD-biplot may identify more meaningful taxa with respect to the separation of the samples than the AMD and SVD biplots.

### Analysis of human gut microbiome data

We consider the human gut microbiome data collected in a study of healthy children and adults from the Amazonas of Venezuela, rural Malawi and US metropolitan areas [15]. The original data set **X** consists of counts for 149 taxa for 100 samples. The squared unweighted UniFrac distance matrix **Δ** ∈ ℝ^100×100^, computed using the `R` package `phyloseq` [16], was used to measure the distance between samples. Here, the distance between two samples is based entirely on the number of branches they share on a phylogenetic tree. The distance hence accounts only for the presence/absence of each taxon (not its abundance). The corresponding similarity kernel **H** was then derived as 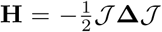, which is a positive semi-definite matrix with rank 99. Let 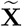 and 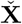, respectively, denote the CLR transformed data and the relative abundance data. Similar to the first study, the GMD-biplot and the AMD-biplot were constructed based on the similarity kernel **H** using 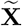 and 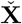 respectively, and the SVD-biplot was constructed based on the SVD of 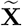.

As concluded in [15], shared features of the functional maturation of the gut microbiome are identified during the first three years of life. We thus define a binary outcome *h*_*i*_ based on the age of each sample as:

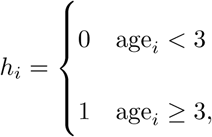

for *i* = 1, …, 100. Approximately 70% of the samples are assigned to group 0 and the remaining 30% are assigned to group 1.

In all biplots, the *i*^*th*^ sample is colored by age_*i*_ and symbolized by *h*_*i*_. Fig. 2(d) indicates that more than 80% of the total variance is explained by the GMD-biplot in Fig. 2(a), which provides a good visualization of sample clusters across age. By examining the relationship between the arrows and the color of the sample points in Fig. 2(a), we see that *prevotella* may be elevated in adults, while *parabacteroides* appears to be elevated in infants. In contrast, Fig. 2(e) shows that less than 15% of the total variance is explained by the AMD-biplot in Fig. 2(b) and the AMD values are not decreasing. As shown in Fig. 2(b), the AMD-biplot also displays potential clusters across age, but the sample points are not as tightly clustered as those in Fig. 2(a). *Odoribacter* appears to be elevated in adults in Fig. 2(b), while *lactobacillus* appears associated with infants. As a reference, Fig. 2(c) shows the SVD-biplot of 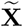, which looks very similar to Fig. 2(a).

To further quantify the classification accuracy, for each biplot we predicted the probability that each sample belongs to group 1 based on leave-one-out cross validation using the binary logistic regression of the group index *h*_*i*_ on the two selected components. We then plotted an ROC curve for each biplot based on the predicted probabilities (Fig. S1) and calculated the area under the ROC curve (AUC): the GMD, AMD and SVD biplots, respectively, yield an AUC of 0.989, 0.976 and 0.990. The AUC results indicate that the GMD-biplot provides a better separation of age groups than the AMD-biplot, but there is not a clear difference between the GMD-biplot and the SVD-biplot. This may be because, for the CLR-transformed data 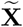, the unweighted UniFrac distance is not as informative with respect to age as the weighted UniFrac distance was in the tobacco data with respect to product groups.

We emphasize that both the GMD-biplot and the SVD-biplot identify *prevotella* and *parabacteroides* as top taxa, while the AMD-biplot identifies completely different ones. As [15] confirms that the trade-off between *prevotella* and *bacteroids* (including *parabacteroides*) considerably drives the variation of microbiome abundance in adults and children between 0.6 and 1 year of age in all studied populations, the GMD and SVD biplots may thus identify more biologically meaningful taxa than the AMD-biplot. It should, however, be noted that these bacterial are “identified” based on circumstantial, not statistical, evidence, and more work is needed to examine statistical associations in this context.

### Incorporating a kernel for variables into the GMD-biplot

The GMD problem defined in Eq. (1) allows not only the similarity kernel for samples, but also a kernel for the variables. Including both kernels may further improve the accuracy of sample classification as well as the identification of important variables. We illustrate this advantage by incorporating a kernel for variables in the analysis of the human gut microbiome data. More specifically, we first calculate a matrix **Δ**_*R*_ ∈ ℝ^149×149^ of squared patristic distances between the tips of the phylogenetic tree for each pair of taxa and then derive a similarity matrix **R** as 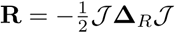. Fig. 3(a) shows the GMD-biplot with the additional kernel **R** incorporated. The ROC analysis based on the leave-one-out cross validation for Fig. 3(a) yields an AUC of 0.984, which is higher than that of the AMD-biplot (Fig. 2(b)) but slightly lower than Fig. 2(a) and Fig. 2(c). This may be because both **H** and **R** highly depend on the phylogenetic tree. Thus, incorporating **R** may be redundant and may reduce the accuracy of the sample clustering in this case. The top 3 taxa identified in Fig. 3(a) include *prevotella* but not *parabacteroides*, which may explain the lower clustering accuracy.

Including an additional kernel for variables in the GMD-biplot is related to method of double principal coordinate analysis (DPCoA) [17]. DPCoA, as shown in [18], is equivalent to a generalized PCA which essentially incorporates an additional similarity kernel for variables into the analysis, as described in Proposition 1, but for **H** = **I**_*n*_. As suggested in [19], DPCoA can provide a biplot representation of both samples and meaningful taxonomic categories. Hence, the GMD-biplot can also be viewed as an extension of DPCoA biplots because the GMD allows kernels for both samples and variables, while DPCoA only allows a kernel for variables.

### Simulation

In this section, we conduct a simulation study based on the smokeless tobacco data to illustrate a scenario in which the AMD-biplot may fail to separate the samples, whereas the GMD-biplot performs well. Let **H**^*^ and **X** be the similarity kernel and data matrix from the smokeless tobacco data, respectively. We consider the eigen-decomposition of **H**^*^ as **H**^*^ = **BΛB**^*T*^ : **B** is a 45 × 27 matrix whose columns are eigenvectors of **H**^*^ and **Λ** = diag(*λ*_1_, …, *λ*_27_) is a diagonal matrix whose elements are the eigenvalue of **H**^*^. Then, the AMD-biplot is based on the following approximated orthogonal decomposition of **X**:

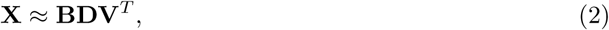

where **D** = diag(*d*_1_, …, *d*_27_) and **V** is a 271 × 27 matrix with orthonormal columns. As shown in Fig. 2(d), *d*_1_, …, *d*_27_ may not be decreasing. For *j* = 1, …, 27, we define

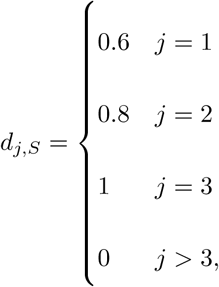

and construct the simulated data set **X**_*S*_ as **X**_*S*_ = **BD**_*S*_ **V**^*T*^, where **D**_*S*_ = diag(*d*_1,*S*_, …, *d*_27,*S*_). For *i* = 1, …, 45, we define a binary outcome *w*_*i*_ that indicates the group index of the *i*^*th*^ sample as:

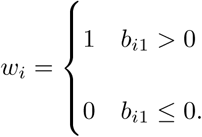

The GMD-biplot and the AMD-biplot of **X**_*S*_ with similarity measure **H**^*^ are presented in Fig. 4(a) and 4(b), respectively. It can be seen that the two groups are completely mixed up in the AMD-biplot because the first column of **B** is not selected for visualization. In contrast, the GMD-biplot successfully visualizes the sample groups by displaying the first and second GMD components.

**Figure 4:**
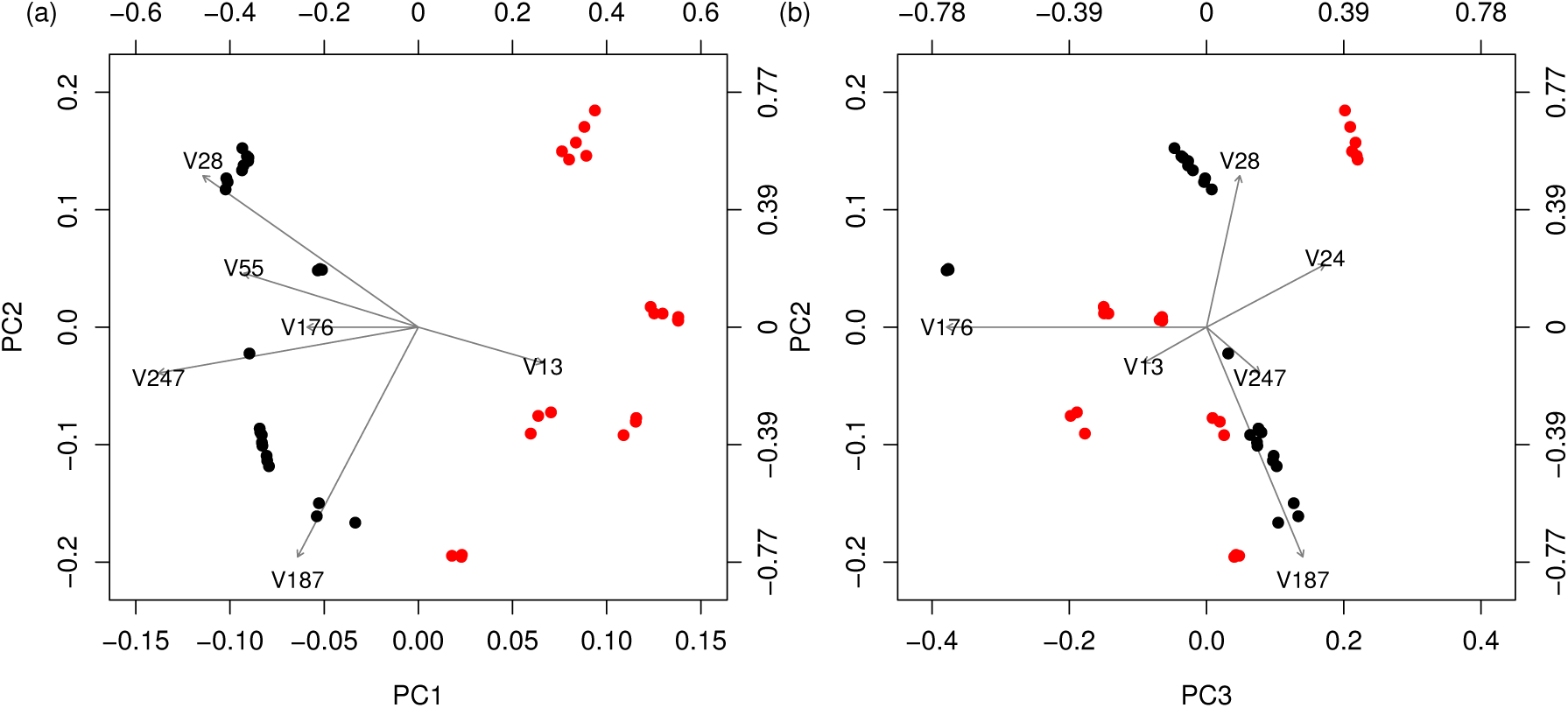
Biplots for the analysis of the simulated data. (a): The GMD-biplot based on the first and second GMD components; (b): The AMD-biplot based on the second the third components; both biplots display the top 6 taxa with the longest arrows. The samples points are colored by the group index (1 = “red”; 0 = “black”).

To see why this occurs, we summarize the first three diagonal elements of **Λ, D**_*S*_ and 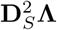 in Table 1 and notice that *d*_1,*S*_ *< d*_2,*S*_ *< d*_3,*S*_. Consequently, the AMD-biplot displays the second and third columns of **BD**_*S*_, and hence it completely fails to classify the samples because the group index *w*_*i*_ only depends on the first column of **B**. In contrast, Proposition 1(a) shows that the GMD-biplot is based on the two largest eigenvalues and the corresponding eigenvectors of 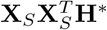. It can be further seen that

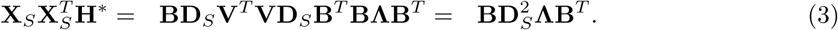

**Table 1:** The first three diagonal elements of Λ, D_*S*_ and 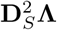 in the simulation.

|  | 1<sup>st</sup> | 2<sup>nd</sup> | 3<sup>rd</sup> |
| --- | --- | --- | --- |
| $\mathbf{D}_S$ | 0.6 | 0.8 | 1 |
| $\mathbf{\Lambda}$ | 3.09 | 1.26 | 0.77 |
| $\mathbf{D}_S^2\mathbf{\Lambda}$ | 1.11 | 0.81 | 0.77 |

Eq. (3) implies that the diagonal elements of 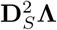 are the eigenvalues of 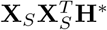 and columns of **B** are the corresponding eigenvectors. Hence, it can be seen from Table 1 that 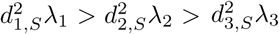, even though *d*_1,*S*_ *< d*_2,*S*_ *< d*_3,*S*_. Therefore, the GMD-biplot displays the first and second column of **BD**_*S*_ **Λ**^1*/*2^ as sample points, which successfully captures sample classifications.

## Discussion

Biplots have gained popularity in the exploratory analysis of high-dimensional microbiome data. The traditional SVD-biplot is based on Euclidean distances between samples and cannot be directly applied when more general dissimilarities are used. Since Euclidean distances may not lead to an optimal low-dimensional representation of the samples, we have extended the concept of the SVD-biplot to allow for more general similarity kernels. The phylogenetically informed UniFrac distance, used in our examples, defines one such kernel. In settings where a general (possibly nonlinear) distance matrix is appropriate, our approach provides a mathematically rigorous and computationally efficient method, based on the GMD, that allows for plotting both the samples and variables with respect to the same coordinate system.

Our first data example with the smokeless tobacco data set from [11] demonstrates the merits of the proposed GMD-biplot. We found that the GMD-biplot successfully displays different types of products, while the AMD-biplot is not able to completely separate dry and moist snuffs and the SVD-biplot fails to separate moist and toombak samples. As shown in Table S1, the GMD-biplot is also able to identify biologically more meaningful taxa that are related to the different types of products, compared to the AMD-biplot and the SVD-biplot.

In our second example, the GMD-biplot also outperforms the AMD-biplot in terms of both the sample clustering and the identification of important taxa. However, there is not a clear advantage of the GMD-biplot over the SVD-biplot in this example. This difference between the two examples may be attributed to the relation between the Euclidean kernel and the non-Euclidean similarity measure. Denoting by **XX**^*T*^ and **H** the Euclidean kernel and the non-Euclidean similarity measure, respectively, it can be seen that the sample configuration in the AMD-biplot and the SVD-biplot depend solely on either **H** or **XX**^*T*^, whereas the GMD-biplot uses the top two eigenvectors of **XX**^*T*^ **H**, the matrix product of the Euclidean kernel **XX**^*T*^ and **H**. Hence, if **XX**^*T*^ contains substantially more information about sample clustering than **H**, then taking **H**^*T*^ into consideration may not further improve the accuracy of sample clustering. Indeed, this may be the case in our second example, where the clustering of samples using the Euclidean distance between samples of the CLR transformed data is highly successful because the difference of the microbial profiles between infants and adults is obvious even without the help of the UniFrac distance. However, a possibly more common scenario is when both **H** and **XX**^*T*^ contain some, but different, information on sample clustering. In such cases, taking both **XX**^*T*^ and **H** into consideration may improve the sample clustering and provide better biological interpretation.

In practice, we typically do not know what the true configuration of samples look like, so it is impossible to determine whether **H** or **XX**^*T*^ contains more information about sample clusters. Also, it is sensible to assume that **XX**^*T*^ and **H** are “co-informative” in the sense that they exhibit a shared eigenstructure; for instance, both may be informative for clustering samples. The co-informativeness can be quantified precisely using the Hilbert-Schmidt information criteria (HSIC) [20]. For any two kernels **K**_1_ and **K**_2_, the empirical HSIC is proportional to tr(**K**_1_**K**_2_). Hence, by definition, the GMD problem (1) is equivalent to minimizing the HSIC between (**X** − **USV**^*T*^) (**X** − **USV**^*T*^)^*T*^ and **H** over **U, S** and **V**. In other words, if we consider **X** − **USV**^*T*^ as the residual matrix of **X**, then the GMD solutions can be interpreted as the best approximation to **X** in the sense that the HSIC between **H** and the Euclidean kernel of the residual matrix is minimized. Thus, the GMD-biplot considers the co-informativeness of **XX**^*T*^ and **H**. Therefore, in many cases it would be a more robust way to display the sample points compared to the AMD-biplot or the SVD-biplot. Another advantage of the GMD-biplot over the AMD-biplot is illustrated in our simulation study. Since the AMD may not give decreasing singular values, the AMD-biplot may not be able to display the most informative eigenvectors of **H**, and may thus fail to cluster the samples. In contrast, the GMD assures that the resulting singular values are non-increasing.

Our discussion in this paper has focused on the *form biplot*, which aims to visualize the relationship between variables and the sample clustering. In other scenarios, where the variation of the data matrix explained by each variable is of particular interest, the *covariance biplot* may be more appropriate. This biplot considers the GMD of **X** with respect to **H**; i.e. **X** = **USV**^*T*^, where **U**^*T*^ **HU** = **I**_*q*_ and **V**^*T*^ **V** = **I**_*q*_. Note that

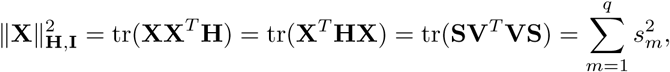

where **S** = diag(*s*_1_, …, *s*_*q*_). Furthermore, since **V** has orthogonal columns, it can be seen that 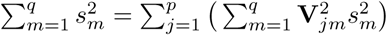. Thus, the value of 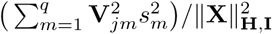 gives the proportion of the variability in 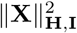 explained by the *j*^*th*^ variable. Note that when *q* = 2, the length of the arrow of the *j*^*th*^ variable in the covariance biplot is given by 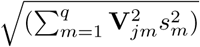. Therefore, in a covariance biplot, the arrows shed light on how the total variance of the data is partitioned into parts explained by each variable.

## Supplemental Material

**Table S1** Ranks of the top 10 taxa identified by the GMD, AMD and SVD biplot in the analysis of smoke tobacco data.

**Fig S1** ROC curves for the GMD, AMD and SVD biplots in the analysis of human gut microbiome data.

## Acknowledgements

This work was partially supported by grant R01 GM114029 from the NIH. AS also acknowledges the support from the NSF through grant DMS-1561814. JM and TR acknowledge support by grant R01 GM129512 from the NIH. TR also acknowledges support from NIH grants R01 CA192222 and P01 CA168530. The content of the manuscript is solely our responsibility and does not necessarily represent the official views of the NIH or any other funding agency. We are thankful to Parker Knight for his assistance with the software development.

## Data Availability

All data used are publicly available in [11] and [15]. All computations are carried out in the `R` programming language and the proposed biplot is implemented in our `R` package “GMDecomp”, available at “https://github.com/taryue/GMDecomp“.

