## Supplementary material for "The GMD-biplot and its application to microbiome data": Table_S1

mSystems Submission Template mSystems Submission Template mSystems Submission Template mSystems Submission Template mSystems Submission Template mSystems Submission Template

| GMD | | AMD | | SVD | |
|---|---|---|---|---|---|
| taxa | rank | taxa | rank | taxa | rank |
| halophilus | 26 | succinus | 137 | succinus | 137 |
| alloiococcus | 22 | lactobacillus | 67 | aerococcaceae3 | 134 |
| aerococcaceae1 | 64 | aerococcaceae3 | 134 | corynebacterium | 50 |
| tetragenococcus | 33 | corynebacterium | 50 | staphylococcus | 139 |
| aerococcaceae3 | 134 | stationis | 15 | alloiococcus | 22 |
| stationis | 15 | enterobacteriaceae | 12 | enterobacteriaceae | 12 |
| enterobacteriaceae | 12 | halophilus | 26 | staphylococcus | 161 |
| aerococcaceae2 | 42 | alloiococcus | 22 | halophilus | 26 |
| yaniella | 24 | acetobacter | 131 | aerococcaceae1 | 64 |
| granulicatella | 48 | yaniella | 24 | enterobacteriaceae | 17 |
| avg. rank | 42 | avg. rank | 61.8 | avg. rank | 76.2 |
