## Supplementary figures and images for "The GMD-biplot and its application to microbiome data"

### Fig_S1

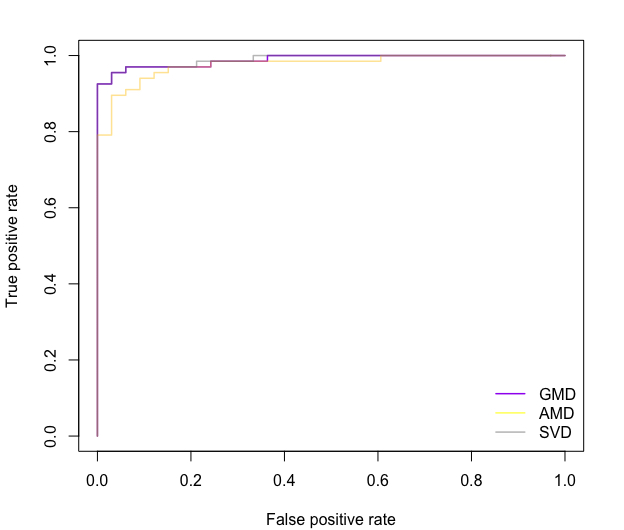
